# Evaluation of Machine Learning Models for Aqueous Solubility Prediction in Drug Discovery

**DOI:** 10.1101/2024.06.10.598383

**Authors:** Nian Xue, Yuzhu Zhang, Sensen Liu

## Abstract

Determining the aqueous solubility of the chemical compound is of great importance *in-silico* drug discovery. However, correctly and rapidly predicting the aqueous solubility remains a challenging task. This paper explores and evaluates the predictability of multiple machine learning models in the aqueous solubility of compounds. Specifically, we apply a series of machine learning algorithms, including Random Forest, XG-Boost, LightGBM, and CatBoost, on a well-established aqueous solubility dataset (i. e., the Huuskonen dataset) of over 1200 compounds. Experimental results show that even traditional machine learning algorithms can achieve satisfactory performance with high accuracy. In addition, our investigation goes beyond mere prediction accuracy, delving into the interpretability of models to identify key features and understand the molecular properties that influence the predicted outcomes. This study sheds light on the ability to use machine learning approaches to predict compound solubility, significantly shortening the time that researchers spend on new drug discovery.

## I. Introduction

Aqueous solubility, a key determinant of drug absorption, distribution, metabolism, and excretion (ADME), significantly impacts drug efficacy and bioavailability [1]. Limited aqueous solubility (often expressed as *logS*) can significantly hinder a drug candidate’s effectiveness by restricting its bioavailability [2]. Therefore, the early identification of molecules with unfavorable aqueous solubility during the drug discovery pipeline is critical for reducing the risk of failure and expediting the development of successful therapeutics [3].

While experimental solubility measurements are resource-intensive and time-consuming, *in-silico* prediction methods offer a promising alternative [4]. By harnessing the wealth of information encoded within molecular structures, these methods offer a powerful approach to estimating a compound’s aqueous solubility without the need for laboratory experiments [5]. However, the accurate and rapid prediction of aqueous solubility has posed a significant challenge in drug discovery for decades. Computational methods, particularly artificial intelligence (AI), offer promising avenues for addressing these challenges [6]. In the context of aqueous solubility prediction, this translates to utilizing a large set of molecules with known solubility values (training data) to establish a relationship between a molecule’s feature (represented by molecular descriptors) and its corresponding *logS* value. After training, the model can quickly predict the *logS* values of new molecules. This enables rapid screening and prioritization of drug candidates with favorable solubility profiles, accelerating drug development.

So far, a variety of ML algorithms have emerged as powerful tools for predicting aqueous solubility. However, they have their own strengths and limitations. For example, Support Vector Machines (SVMs) excel with smaller datasets but may struggle with larger ones [7]. While Random Forest (RF) models and tree-based boosting methods like LightGBM have been investigated, studies suggest superior generalization performance with RF [8]. Artificial Neural Networks (ANNs), while powerful, can be prone to overfitting and noise sensitivity [9]. Deep learning offers improved accuracy through automatic feature extraction [10]– but may require extensive training data and sacrifice interpretability [13]. Ultimately, the choice of algorithm depends on dataset size, desired accuracy, and interpretability needs.

However, ML model selection and optimization for solubility prediction remain challenging, necessitating careful consideration of feature selection, model complexity, and data quality. In addition, despite successful application of numerous ML algorithms, a comprehensive comparison of their performance on a common dataset, with emphasis on interpretability and feature importance, is lacking. This study aims to fill this gap by comparatively analyzing prominent ML algorithms, including Random Forest, XGBoost, LightGBM, and CatBoost.

This paper explores and evaluates ML models for aqueous solubility prediction in drug discovery. Using the Huuskonen dataset [14], we train and optimize models via hyperparameter tuning, then investigate their interpretability through feature importance analysis. SHapley Additive exPlanations (SHAP) analysis [15] is employed to further reveal key features influencing solubility. Our comprehensive analysis provides insights into the strengths and weaknesses of different ML approaches, guiding researchers in selecting the most suitable algorithm based on dataset size, desired accuracy, and interpretability needs [16].

The major contributions of our paper are summarized as follows:

- Training multiple machine learning models on the Huuskonen dataset for aqueous solubility prediction.
- Investigating the impact of different parameter search strategies on model optimization and prediction performance.
- Analyzing feature importance of four ML models using Shapley values to identify key determinants of solubility.

The remaining of the paper is organized as follows. Section II details the composition of the dataset used for training and evaluating the machine learning models. Section III presents the results, including model performance comparisons, hyperparameter tuning analysis, and feature importance exploration. Section IV discusses the implications and limitations of these findings.

## II. Methods

This section details the dataset used in our experiments, environment setup, ML models employed to predict molecular solubility *logS* and the evaluation metrics used to assess their performance.

### A. Dataset Preparation

In our study, we leveraged the well-established Huuskonen dataset [14], a publicly available benchmark dataset commonly used in literature for solubility prediction tasks. This dataset contains information for 1297 unique organic molecules. Each molecule in the Huuskonen dataset is labeled by its corresponding aqueous solubility value, expressed in units of log mol/L at a temperature range of 20−25°*C* (*pH* = 7.5). Fig. 1 shows the distribution of the value of molecular solubility (logS) in the Huuskonen dataset. Fig.2 shows the visualization of some molecule samples with relatively high solubility in the datasets.

**Fig. 1:**
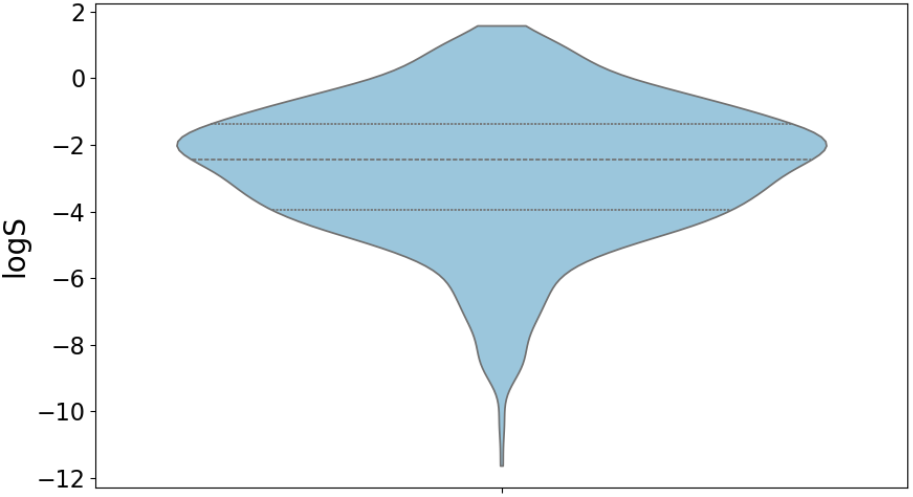
Distribution of the molecular solubility *logS* in the Huuskonen dataset.

**Fig. 2:**
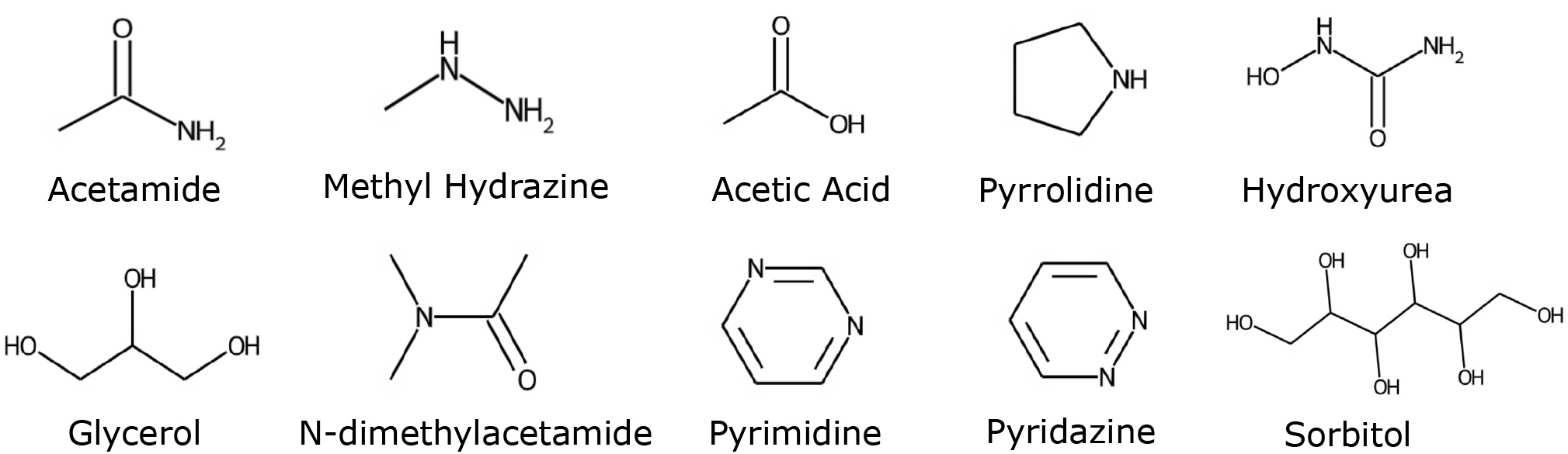
Visualization of representative molecule samples with high solubility.

### B. Experimental Environment Setup

#### Hardware Environments

All ML models were trained and tested on a MacBook Pro equipped with the Apple^®^ M2 chip featuring a 12-core CPU, 19-core GPU, and 16-core Neural Engine. The device also has 32GB of unified memory and enough disk space to store the data and trained models.

#### Software Environments

The MacBook Pro runs a 64-bit macOS system. All the experiments are coded in Python 3.11. Other necessary software packages used in our test include numpy(v1.26.4), pandas(v2.1.4), SHAP(v0.45.0), and scikit-learn(v1.2.2) [17]. Among these packages, numpy and pandas serve as tools for general data processing, scikit-learn is used for building the ML models, while SHAP facilitates explanation of the output generated by ML models.

To prepare the molecular data for ML models, we employed RDKit, a powerful open-source cheminformatics toolkit [18]. RDKit takes the SMILES strings [19], a concise and machine-readable representation of a molecule’s structure, as input. For each molecule in the Huuskonen dataset, RDKit generates a rich set of 210 molecular descriptors. These descriptors serve as the features for our machine learning models and capture various physicochemical characteristics that potentially influence aqueous solubility, such as atom types, bond lengths, and functional groups.

Through data cleaning and filtering processes, the final dataset used for training and testing the models contains 1238 molecules.

### C. Dataset splitting strategy and Machine Learning Models

The dataset was rigorously split into training and test sets following an 80*/*20 ratio using random stratified sampling. This translates to 990 samples allocated for training the models and 248 samples reserved for testing. This split ensures the models are trained on a representative portion of the data while reserving a separate set for unbiased evaluation of their generalizability.

To develop intrinsic solubility models of chemicals based on their structural attributes, we investigated a diverse range of machine learning algorithms, including:

- Linear Regression (LR)
- Random Forest (RF)
- Decision Tree (DT)
- XGBoost (XGB)
- LightGBM (LGBM)
- CatBoost (CatB)

We chose the listed machine learning algorithms because they offer a straightforward interpretation of the relationship between features and target variables, which is crucial for understanding the influence of structural attributes on solubility. In addition, the selected algorithms have demonstrated effectiveness in various domains and are widely used for regression tasks [20]. Finally, compared to models like ANNs, K-Nearest Neighbors, SVM, and Naive Bayes, the chosen algorithms typically require less computational resources and are easier to implement, making them practical choices for experimentation and deployment [21].

### D. Evaluation Metrics

Following training, the performance of each model is evaluated by two core metrics: R-squared (*R*^2^) and Mean Squared Error (*MSE*). In machine learning, *R*^2^ serves as a prevalent metric for evaluating the linear relationship between true and predicted values, while *MSE* are commonly employed as performance metrics, which are defined as follows:

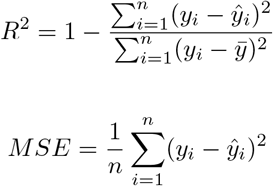

where *n* is the number of observations, *y*_*i*_ represents the observed values, *ŷ*_*i*_ represents the predicted values, and 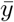 represents the mean of the observed values.

These metrics were calculated on both the training and the test datasets. R-squared reflects the proportion of variance in the target variable (*logS*) explained by the model, while *MSE* measures the average squared difference between the predicted and actual *logS* values.

In addition, SHAP analysis [15] is utilized to interpret the predictions generated by the machine learning models, thus facilitating the understanding of the key features influencing outcomes. This approach has garnered considerable attention within the machine learning area owing to its interpretive capacity. Specifically, SHAP provides a way to assign an importance value to each feature based on its shapley value and estimates the contribution of each feature to the final output. The equation can be represented as:

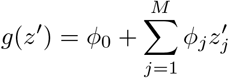

where *g* is the model, *M* is the number of input features, *z*^*′*^ indicates whether the corresponding feature exists (1, 0), *ϕ* is the attribution value (i. e., Shapley value) of each feature, and *ϕ*_0_ is a constant.

## III Results

In this section, we present a comprehensive evaluation of the performance of the ML models and hyperparameter optimization for various machine learning models used to predict molecular solubility, *logS*. As a measure of a compound’s ability to dissolve in a given solvent, typically water, understanding the factors that lead to optimal solubility *logS* is a critical prerequisite in fields such as pharmaceuticals and environmental science.

### A. Model Performance Comparison

Table I summarizes the *R*^2^ and *MSE* scores achieved by each model on both the training and test datasets. Among all the models, the average *R*^2^ and *MSE* scores on testing data are 0.403 and 0.904, and the median *R*^2^ and *MSE* scores on testing data are 0.390 and 0.905, respectively. An initial analysis of the R-squared scores suggests comparable performance across all models, indicating a good overall fit to the data. However, a closer examination of the test set *MSE* scores reveals key differences in generalizability. Tree-based models, particularly CatBoost (0.34 *MSE*), LightGBM, and the fine-tuned Random Forest, obviously outperform other models in terms of predicting unseen data. This is corroborated by the significantly lower *MSE* values achieved on the test set compared to the training set for the default Decision Tree and Random Forest models. These latter observations are indicative of overfitting, where the models prioritize memorizing the training data at the expense of generalizability.

**TABLE I:** Performance Comparison of Machine Learning Models for Solubility Prediction.

| Model | Training Set $MSE$ | Training Set $R^2$ | Test Set $MSE$ | Test Set $R^2$ |
| --- | --- | --- | --- | --- |
| LinearRegression | **0.24** | **0.94** | 0.41 | 0.90 |
| LassoRegression-Normalization | 0.27 | 0.94 | 0.40 | 0.90 |
| LassoRegression-MinMax Scaler | 0.33 | 0.92 | 0.38 | 0.91 |
| DecisionTree (default setting) | 0.46 | 0.89 | 0.64 | 0.84 |
| DecisionTree (tree size = 40 nodes) | 0.30 | 0.93 | 0.60 | 0.86 |
| RandomForest (default setting) | 0.42 | 0.90 | 0.42 | 0.90 |
| RandomForest (ensemble size = 500 trees) | 0.43 | 0.90 | 0.41 | 0.90 |
| RandomForest (mtry = 46) | 0.35 | 0.92 | 0.35 | 0.92 |
| LightGBM | 0.27 | 0.93 | 0.35 | 0.92 |
| CatBoost | 0.26 | 0.93 | **0.34** | **0.92** |
| XGBoost | 0.31 | 0.92 | 0.37 | 0.91 |
| Average | 0.319 | 0.923 | 0.403 | 0.904 |
| Median | 0.305 | 0.925 | 0.390 | 0.905 |

### B. Hyperparameter Tuning for Tree-Based Models

Unlike linear models, tree-based models rely on hyperparameters to manage their complexity. Inappropriate selection of these hyperparameters can lead to overfitting. As evidenced in Table I, the default DT and RF models exhibit this behavior, achieving lower *MSE* on the training set compared to the test set.

To gain a deeper understanding of how hyperparameters influence model performance, we conducted a grid search with 3-fold cross-validation (CV) for DT and RF models. This technique systematically evaluates a range of hyperparameter values and assesses their impact on chosen metrics, (i.e., *R*^2^ and *MSE*).

#### 1) Decision Tree (DT)

We conducted a grid search to explore the impact of tree size, controlled by the max leaf nodes parameter, on the Decision Tree model’s performance. As evidenced by Table I, the fine-tuned model (row 5) achieves superior performance on the test set compared to the default model (row 4), which suffers from overfitting. Fig. 3 visually depicts the key findings from this grid search analysis.

The left panel of Fig. 3 reveals a divergence in *R*^2^ scores: training data *R*^2^ increases until plateauing, while test data *R*^2^ plateaus earlier (around 40 nodes). This suggests overfitting with larger trees. Mirroring this, the right panel shows a continuous decrease of *MSE* on the training set with increasing tree size, but the *MSE* on the test set remains flat after exceeding 40 nodes.

**Fig. 3:**
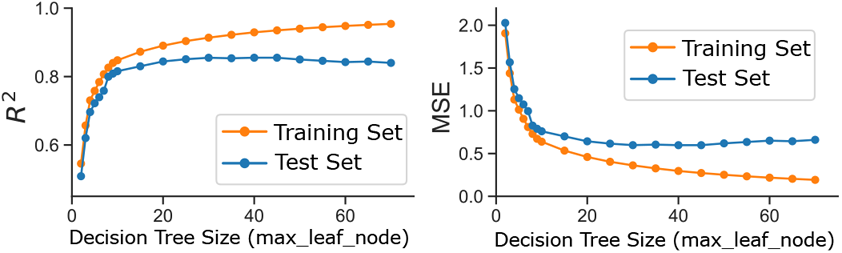
Grid search reveals the impact of tree size on Decision Tree performance. Left: Decision Tree *R*^2^ vs. tree size (training and test). Right: Decision Tree *MSE* vs. tree size (training and test).

#### 2) Random Forest

While the default RF parameters yield comparable test set performance, an analysis of the out-of-bag (OOB) data revealed signs of over-fitting in the training set. To verify this, we further conducted a grid search focusing on the ensemble size (number of trees in the forest). As depicted in Fig. 4, the grid search explored a range of ensemble sizes and evaluated their impact on both OOB *R*^2^ and *MSE*.

Fig. 4 suggests that increasing the ensemble size leads to a gradual improvement in OOB *R*^2^ and a decrease in OOB *MSE*, indicating enhanced model performance for aqueous solubility prediction. Notably, this improvement plateaus around an ensemble size of 500 trees. Therefore, based on the observed trends and the trade-off between computational cost and performance, an ensemble size of 500 was identified as the optimal for aqueous solubility prediction.

**Fig. 4:**
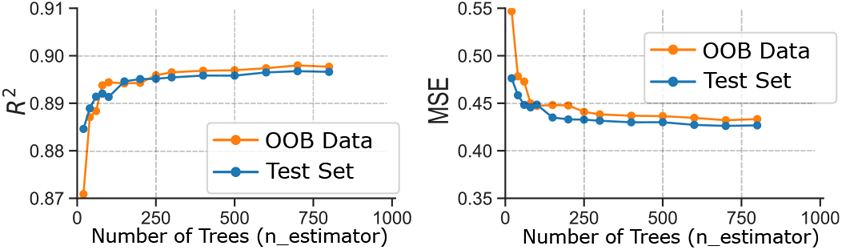
Grid search exploration of ensemble size for the Random Forest model. Left: *R*^2^ score on the training (OOB) and test data as a function of ensemble size (number of trees). Right: *MSE* score on the training (OOB) and test data as a function of ensemble size(number of trees).

#### 3) Gradient Boosting Algorithms

Gradient boosting algorithms (LightGBM, CatBoost, XGBoost) were hyperparameter tuned using randomized search cross-validation. We set the number of sampled parameter combinations to 200 (*n iter* = 200) and used the default *R*^2^ score as the objective function. In addition, we implemented 3-fold cross-validation, where each data fold is used for validation once to estimate generalization error. The parameter search results are summarized in Table II.

Hyperparameter tuning revealed a trade-off between model complexity, performance, and training/inference times. We analyze these factors in detail below.

**TABLE II:**
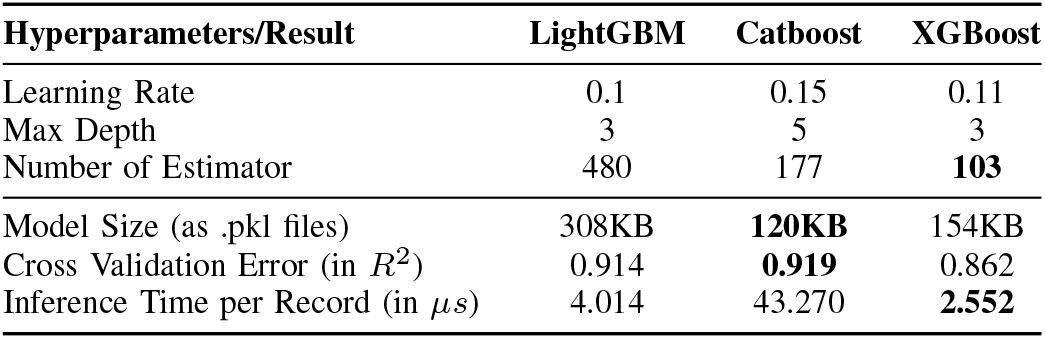
Hyperparameter Tuning for Gradient Boosting Algorithms.

#### Model File Size and Complexity

The trained models were saved as “pkl” files, and their file sizes were compared. LightGBM produced the largest file size (308 KB), likely due to its higher number of estimators identified during the CV process. Interestingly, XGBoost (154 KB) was slightly larger than CatBoost (120 KB), despite lower model complexity based on *R*^2^ score, suggesting potential differences in model architecture or data representation.

#### Cross-Validation Performance

Cross-validation reveals that CatBoost has the highest *R*^2^ score, indicating a potentially superior overall fit among the three models. This aligns with its smaller file size, suggesting a more compact and efficient model representation. LightGBM achieved comparable performance, while XGBoost had the lowest *R*^2^ score. However, it is important to consider the trade-offs involved.

#### Training and Inference Time

While individual prediction inference times were negligible for all models (in *µs*), CatBoost exhibited the slowest inference, roughly 15 times longer than XGBoost. This may be insignificant for small datasets but can become noticeable for larger ones. LightGBM and XGBoost offer a good balance between accuracy and speed, making them suitable for computationally demanding scenarios.

Also, our findings on the performance trade-offs between these models are corroborated by existing literature ([22]). Studies have consistently shown that LightGBM prioritizes fast training speeds, while CatBoost offers improved accuracy at the expense of longer training times.

## IV Discussion

### A. Insights from Solubility Predictions and Significance

To illustrate the predictive capabilities of our machine learning approach, we focus on the CatBoost model, which achieved the highest performance among the five models evaluated (Table I). Table III displays the solubility predictions for the top 5 most soluble and top 5 least soluble drug molecules identified by the CatBoost model.

**TABLE III:** Analysis of Top Predicted Soluble and Insoluble Compounds.

| Molecule | Solubility ($\log S$) | Uses |
| --- | --- | --- |
| 2,2',3,3',5,5',6,6'-PCB | -9.80 | Industrial Coolant |
| 2,3,4,5,6-PCB | -8.25 | Industrial Coolant |
| 2,2',3,4,5,5'-PCB | -7.78 | Industrial Lubricant |
| o,p'-DDT | -7.69 | Pesticide |
| Naphthacene | -7.43 | Organic Diode |
| Piperazine | 0.51 | Anthelmintic Agent |
| N-methylpyrrolidone | 0.56 | Solvent |
| O-ethyl carbamate | 0.68 | Anesthetic Agent |
| Succinic Acid | 0.70 | Curative Agent |
| N-methylurea | 0.73 | Antitumor Agent |

These two sets of molecules hold particular significance for drug discovery. Highly soluble compounds are desirable as they readily dissolve in aqueous solutions, facilitating oral administration and promoting bioavailability. Conversely, poorly soluble drugs can present formulation challenges and potentially hinder their effectiveness. Therefore, analyzing both ends of the solubility spectrum provides valuable insights for optimizing drug design and development, especially considering the critical role water solubility plays in pharmacokinetics and pharmacodynamics.

#### 1) Machine Learning at the Forefront of Drug Discovery

The ability to accurately predict solubility using machine learning models like CatBoost represents a significant advancement in the field of drug discovery. Traditionally, solubility determination relied on laborious experimental techniques. ML offers a powerful alternative, enabling rapid *in-silico* assessment of a compound’s solubility profile. This translates to substantial time and cost savings during the early stages of drug development. Furthermore, by identifying highly soluble drug candidates early on, ML can expedite the development of orally administered medications, a preferred route for patient convenience and improved treatment adherence. In essence, ML empowered solubility prediction is poised to become a cornerstone of efficient and cost-effective drug discovery pipelines within the pharmaceutical industry.

#### 2) Analysis of Least Soluble Molecules

Intriguingly, the CatBoost model identified a cluster of three structurally related molecules within the top five predicted least soluble compounds. These belong to the polychlorinated biphenyl (PCB) group. PCBs’ low aqueous solubility, stemming from their chemical stability, leads to environmental persistence, bioaccumulation, and potential adverse health effects. Cat-Boost’s accurate prediction aligns with established scientific understanding and highlights its potential for environmental applications, such as identifying and monitoring PCB contamination in soil and sediment.

The remaining two least soluble compounds, o,p’-DDT and naphthacene, also exhibit low water solubility. This property contributed to o,p’-DDT environmental persistence and subsequent worldwide ban. While naphthacene, a polycyclic aromatic hydrocarbon, is not a common drug, its low solubility is relevant due to its presence in coal tar and cigarette smoke, both linked to health problems.

It is important to distinguish that, although not therapeutic drugs, PCBs, o,p’-DDT and naphthacene serve as valuable examples in drug discovery. Their low solubility highlights the challenges posed by this property during formulation and *in vivo* effectiveness, emphasizing its importance throughout the drug development process.

#### 3) Analysis of Most Soluble Molecules

The five predicted most soluble molecules encompass a diverse range of organic compounds, including N-methylurea, succinic acid, O-ethyl carbamate, N-methylpyrrolidone, and piperazine. High solubility translates to efficient drug delivery through oral administration, a major advantage for patient compliance. For instance, N-methylurea, a known antitumor agent, benefits from high solubility, enabling the development of oral formulations for improved patient comfort and treatment adher-ence. Similarly, succinic acid, a metabolic intermediate with therapeutic applications, can be readily formulated into oral dosage forms due to its high solubility. Overall, the CatBoost model’s ability to identify highly soluble drug candidates aligns with desirable properties for oral drug delivery in the pharmaceutical industry.

In conclusion, analyzing the solubility predictions for both highly and poorly soluble molecules demonstrates the utility of the prediction model for drug discovery and development. By identifying compounds at opposite ends of the solubility spectrum, the model offers valuable insights for optimizing drug design, formulation strategies, and potential environmental impact. Furthermore, the focus on these top molecules emphasizes the critical role water solubility plays in influencing a drug’s pharmacokinetic and pharmacodynamic behavior.

### B. Feature Importance Analysis

While Section III focuses on optimizing model performance, a subsequent investigation of the underlying data and features used in training is necessary. Given the extensive feature set (200+ descriptors), feature importance analysis allows us to identify molecular features that significantly influence model predictions. By prioritizing these features, we can gain deeper insights into the molecular properties driving solubility within the dataset.

Figure 5 summarizes our results, depicting the top ten features ranked by their importance scores for each of the four machine learning models: RF, LGBM, CatB, and XGB. The x-axis labels display the names of the ten most important features, with the corresponding bar lengths representing normalized importance scores.

**Fig. 5:**
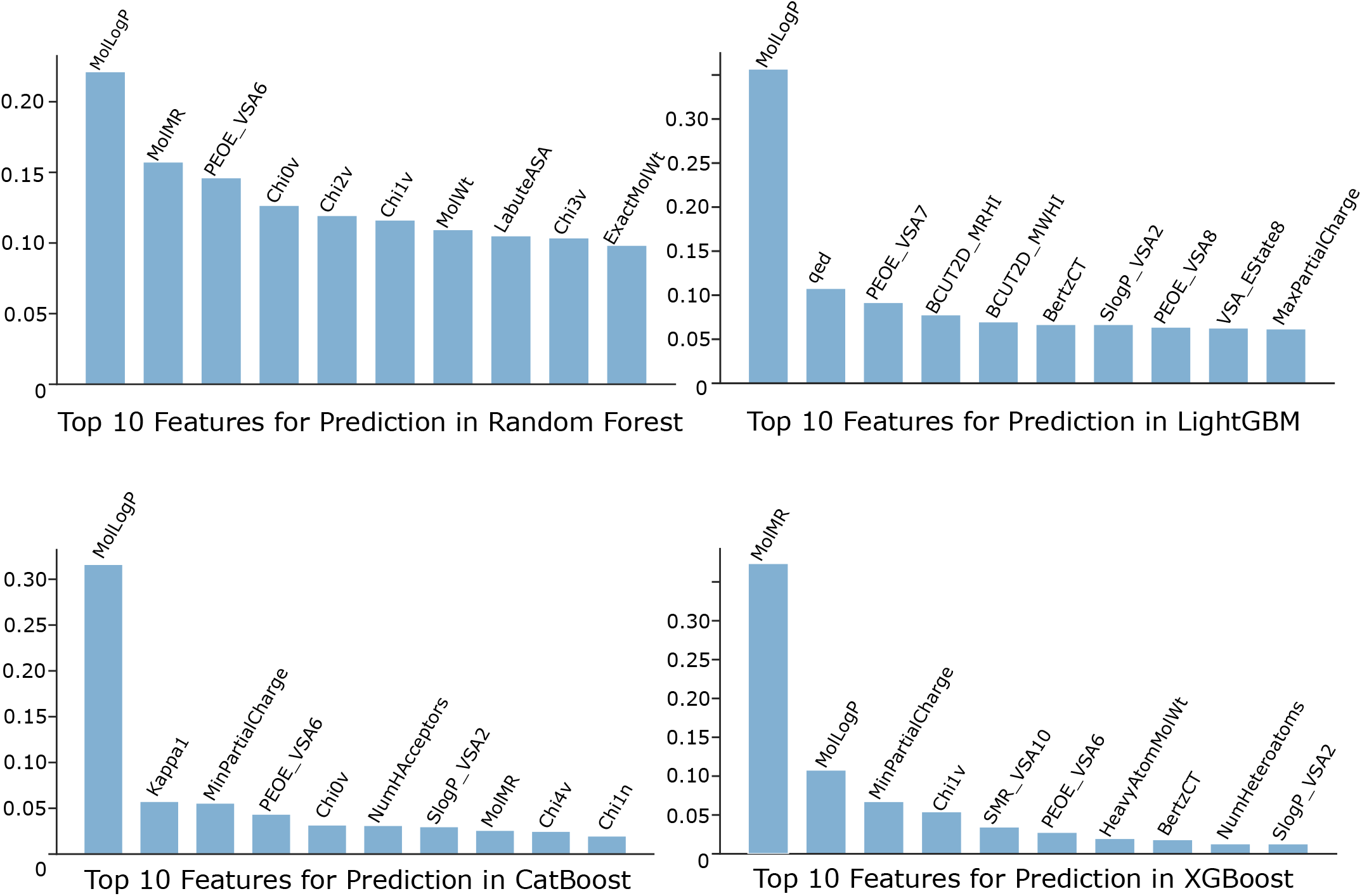
Top 10 features for aqueous solubility prediction across models (RF, LGBM, CatBoost, XGBoost). While molecular hydrophobicity, MolLogP, is dominant for most models, the feature importance distribution in RF suggests a more even contribution from the top features.

A notable observation is that molecular hydrophobicity, represented by MolLogP, exhibits the highest correlation with aqueous solubility, as revealed by three of the models examined (RF, LGBM, and CatBoost in Figure 5). As a descriptor that quantifies a compound’s hydrophobicity, MolLogP value indicates a molecule’s preference for water or organic solvents. In contrast, the XGBoost model stresses the importance of the feature MolMR in determining molecular solubility. MolMR refers to the Wildman-Crippen Molar Refractivity that represents a molecule’s size and polarizability. This finding from the application of XGBoost model suggests that, in addition to the commonly assessed feature of hydrophobicity, molecular size and polarizability may also significantly contribute to aqueous solubility.

Interestingly, the distribution of feature importance scores also provides valuable insights. For instance, the feature importance scores for the RF model exhibit a relatively flat distribution compared to the other models. This suggests that the top ten features in the RF model contribute more equally to the prediction compared to the other models, where the importance scores decrease more sharply for features ranked from 3*rd* to 10*th*.

### C. Deeper Insights from Feature Analysis with SHAP Values

Although the top ten features identified through feature importance analysis, shown in Fig. 5, provide valuable insights into the most influential factors for each model, they offer a limited perspective on guiding drug design. These importance scores represent an ‘average’ influence across the entire training population. To gain a more nuanced understanding of how individual features contribute to predictions for specific data points, we employed SHAP analysis for the CatBoost model, which emerged as the best performing model (Table I).

SHAP values possess two key advantages in this context. First, SHAP values are calculated for every data point to identify potential outliers or features with unusually large influence on specific predictions. Second, SHAP values indicate the direction of a feature’s influence (positive or negative) on the model’s prediction. In contrast, feature importance scores (Fig. 5) only reflect the overall magnitude of a feature’s influence.

Focusing on the most influential feature identified in the CatBoost prediction (Fig. 5), MolLogP, SHAP analysis (Fig. 6) revealed a negative linear-like relationship between MolLogP and its SHAP score. This finding reinforces MolLogP’s importance, as a consistently negative SHAP value distribution across most data points (over 60% with MolLogP values exceeding 2 among the 794 data points) indicates its significant role in the majority of CatBoost model predictions. Furthermore, the negative correlation suggests a potential linear relationship between MolLogP and predicted aqueous solubility within the model.

**Fig. 6:**
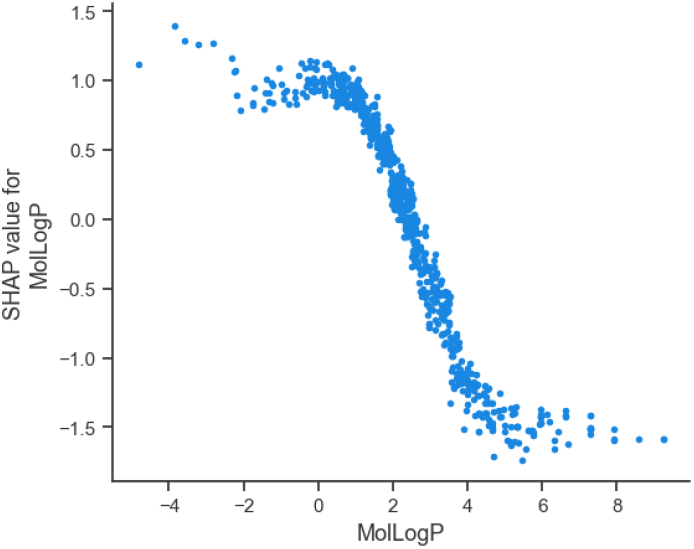
SHAP analysis of MolLogP in the CatBoost model. Hydrophobicity, as a feature, has a strong negative influence on predicted aqueous solubility.

We then extend our investigation to analyze the top 10 most influential features for the CatBoost model using SHAP values. The beeswarm chart in Fig. 7 visualizes these top 10 important features based on their SHAP values, ranked from highest to lowest in terms of importance. The beeswarm interaction plot allows for a more detailed examination of the relationship between these key features and the model’s output. Each dot represents an individual data point’s explanation within a specific feature.

**Fig. 7:**
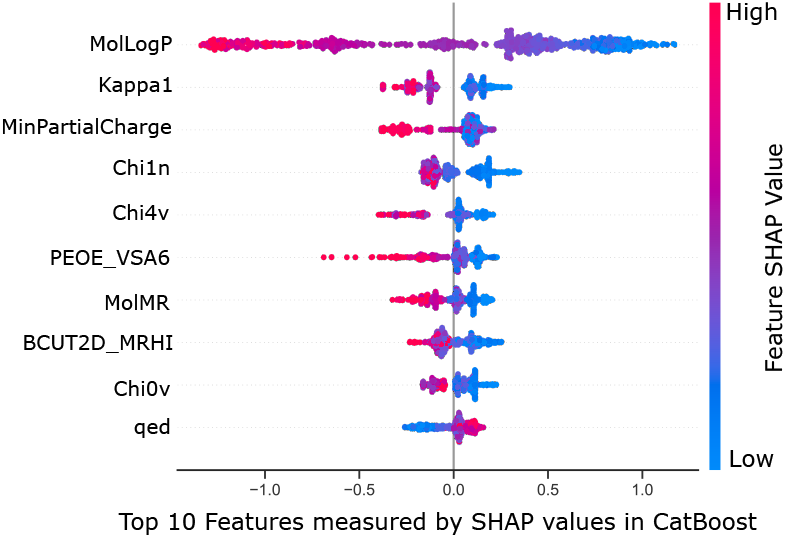
CatBoost model’s SHAP feature importance analysis (top 10 features in beeswarm). While largely consistent with the Feature Importance analysis in Section IV-B, some features differ. The top 3 features, MolLogP, Kappa1, and MinPartialCharge, remain the most important in both analyses.

The x-axis of the beeswarm plot shows the distribution of SHAP values for each key feature, with the color bar linking the feature to the outcome, specifically the solubility value in current study. Distributions centered around zero indicate no impact on the solubility value, while positive or negative centers suggest an increase or decrease, respectively.

#### Consistency with Feature Importance

As expected, MolLogP remains the most important feature according to SHAP analysis, aligning with our initial observations in Section IV-B. In fact, the top three features (“MolLogP”, “Kappa1”, and “MinPartialCharge”) are consistent between both analyses. Other features like “Chi1n”, “Chi4v”, “Chi0v” and “MolMR” also appear in both top 10 lists, albeit with different rankings. However, some discrepancies exist. Features such as “qed”, which stands for quantitative estimation of drug-likeness, [23] and “BCUT2D MRHI”, which refers to the 2D molecular topology, [24], [25] emerge as significant in the SHAP analysis though they were not highlighted in the initial feature importance analysis. Conversely, the initial analysis highlights “SlogP VSA2” and “NumHAcceptors”, which are not reported by SHAP. These findings suggests that SHAP analysis may capture subtle feature interactions that other methods overlook, disclosing important aspects of molecular characterization in predictive modeling.

#### Negative Feature Influence

The color bar indicates the range of negative and positive SHAP values. Interestingly, the top nine most important features appear to be “negative features” based on the clustering of red dots around negative SHAP values (SHAP *<* 0). This suggests that lower values of these features are associated with higher model predictions of aqueous solubility. Unsurprisingly, the feature “qed”, drug-likeness, appears to be a positive feature, with higher values contributing to increased predicted solubility. This correlation, which is not accessible during feature importance score analysis, is logical considering that drug-like molecules often possess properties favorable for solubility in physiological environments.

## V Conclusions

This study explored machine learning for predicting aqueous solubility *in-silico* drug discovery. We have found that Tree-based models, particularly CatBoost and XGBoost, excelled in data prediction (minimal *MSE* on the test dataset). Feature importance analysis further revealed key factors that influence solubility. Although limitations exist, our findings suggest that machine learning holds promise for streamlining drug discovery by identifying molecules with desired aqueous solubility profiles. Future work can explore broader applicability and more complex models for enhanced prediction accuracy, especially when dealing with larger and more complex datasets.

## VI. Acknowledgement

The authors thank Dr. Weiwei He (New York University) for his invaluable contributions, including providing the dataset and initial documents, offering insightful suggestions and engaging in discussions. We are also grateful to Dr. Xinsong Du (Harvard University) for his advice on our manuscript.

